# Perturbed actin cap and nuclear morphology in primary fibroblasts of Huntington’s disease patients as a new phenotypic marker for personalized drug evaluation

**DOI:** 10.1101/2022.04.03.486386

**Authors:** Saja Gharaba, Omri Paz, Lea Feld, Anastasia Abashidze, Maydan Weinrab, Haguy Wolfenson, Miguel Weil

**Author notes:** Corresponding authors Miguel Weil, Haguy Wolfenson.

## Abstract

Human primary skin fibroblast cells from patient’s skin biopsies were used previously as a model to study different neurodegenerative diseases, including Huntington’s Disease (HD). These cells are directly isolated from the patient’s tissue without any alteration in the genome, retaining in culture conditions their endogenous cellular characteristics and biochemical properties, as well as their cellular proliferation capacity for several passages. The aim of this study was to identify a distinctive cellular phenotype in primary skin fibroblasts from various HD patients, using image-based high content analysis, which could be used in the future for personalized drug screening treatment evaluation. We show that HD fibroblasts have a distinctive nuclear morphology associated with a nuclear actin cap deficiency, which in turn affects cell motility in a similar manner to primary skin fibroblasts from Hutchinson-Gilford progeria syndrome (HGPS) patients used as known actin cap deficient cells. Moreover, treatment of the HD cells with either Latrunculin B, used to disrupt actin cap formation, or the antioxidant agent Mitoquinone, used to improve mitochondrial activity, show opposite effects on actin cap associated morphological features and cell motility. The former exacerbates the HD phenotype while the latter improves it. Deep data analysis of the HD nuclear and actin cap features using custom developed image analysis algorithms allow strong cluster classification distinct from HGPS and healthy matching controls, supporting the finding of a novel HD cellular phenotypic marker that could be modulated by pharmacological agents in this patient-based disease model.

## Introduction

Huntington’s disease (HD) is a fatal rare inherited disorder with a broad impact on a person’s functional abilities characterized by unwanted choreatic movements, behavioral and psychiatric disturbances, and dementia^1,2^. HD is an autosomal dominant disease caused by elongated CAG repeats on the short arm of chromosome 4p16.3 in the huntingtin gene (HTT). Normally, the CAG segment is repeated 10 to 35 times within the gene, but in people with HD, the CAG segment is repeated 36 to more than 120 times, causing a mutated translated protein with long Poly glutamine (Poly-Q) repeats called mutant Huntingtin protein (mHtt). The Huntingtin protein is expressed in all human and mammalian cells^3,4^. The role of the Htt protein in humans is unclear. It interacts with over 100 other proteins that are involved in a number of biological functions like transcription, cell signaling, and intracellular transport^5^. Although the mutated protein function is poorly understood, it is toxic to certain cell types due to Poly-Q mediated protein aggregation, particularly in the brain^6^. Notably, it has become clear from different studies that HD is not only a brain disorder, but a multisystem disease that affects numerous cell types^7,8^. It is therefore appealing to use easily accessible cells from patients for phenotype characterization and *in vitro* studies.

Human primary skin fibroblast cells from patient’s skin biopsies were used previously as a model to study different neurodegenerative diseases^9,10^, including HD^11,12^. These cells are directly isolated from the patient’s tissue without any alteration in the genome^13^, retaining in culture conditions their endogenous cellular characteristics and biochemical properties, as well as their cellular proliferation capacity for several passages. This allows performing multiple experiments in a reproducible manner from a single patient’s sample and identify distinctive cellular phenotypes and potential disease biomarkers also expressed in more disease-relevant tissues *in vivo*, as previously described for Amyotrophic Lateral Sclerosis patients’ bone marrow mesenchymal stem cells^14^. Here we adopted image-based high content analysis (HCA) cell phenotyping^15^ as an unbiased approach for screening of robust and distinctive subcellular morphological features in primary skin fibroblasts of HD patients. We found altered nuclear morphology as a significant phenotypic feature of HD cells. Detailed confocal and deep morphometric image analyses of the nuclei, combined with cell migration assays, revealed that this abnormal morphology is due to a nuclear actin cap deficiency in HD patients’ fibroblasts. The nuclear actin cap is an organized apical dome that covers the nucleus and is composed of thick, parallel, and highly contractile actin and myosin filament bundles^16^. Actin cap formation is functionally related to cell migration, intranuclear shaping, chromosomal organization^17^, and cellular mechanoregulation^18^. The actin cap is present in a wide range of adherent eukaryotic cells, but it is disrupted in several human diseases such as cancer^19^ and muscular dystrophy^20^. It is a distinctive biomarker in Hutchinson–Gilford syndrome (HGPS), the accelerated nuclear laminopathy aging syndrome also known as progeria^16,21^. Our results demonstrate that actin cap deficiency is a robust biomarker in HD primary skin fibroblast cells that could be used as a powerful functional measure in drug screening assays as well as for future personalized drug test validations for treatment of HD patients.

## Results

### Primary skin fibroblasts from HD patients display distinct morphological features

To determine phenotypic differences by image-based HCA of primary HD fibroblasts, we analyzed 14 HD and matching human control (HC) cell samples. The primary cells from 5 individuals from each group were plated using a robotic liquid handling unit in 96 well plates in DMEM and 10% FBS and incubated for 24 h at 37°C, 5% CO2, washed, and finally stained with a mix of fluorescent dyes, Hoechst 33342 (blue), calcein-AM (green), mitotracker (far red), and TMRE (red) in HBSS to label the nuclei, cell cytoplasm, mitochondria, and functional mitochondria, respectively. Automated imaging with a 20X objective of thousands of fluorescently stained cells per sample at the four fluorescence channels was performed under environmental controlled conditions using an IN Cell 2200 Analyzer, and image HCA was performed using this system software as described in the Materials and Methods. Figure 1 shows the results from these experiments. Representative micrographs of the labeled cells for image analysis are shown in Figure 1A. Quantitative analysis of these experiments showed clear segregation of the phenotype between the HD and HC cell samples by principal component analysis (PCA) that gave a 64% variance ratio (Figure 1B). The feature histogram distribution by PC dimension indicates a clear difference between the groups (Figure 1C). As shown in Figure 1D, the most relevant HD phenotypic characteristics were mitochondria features and cell morphology, consistent with previous studies^22^. Notably, besides these features, we found that nuclear morphological features are distinctive in the HD cell populations, indicating an HD nuclear phenotype which, to the best of our knowledge, has not been described before.

**Figure 1:**
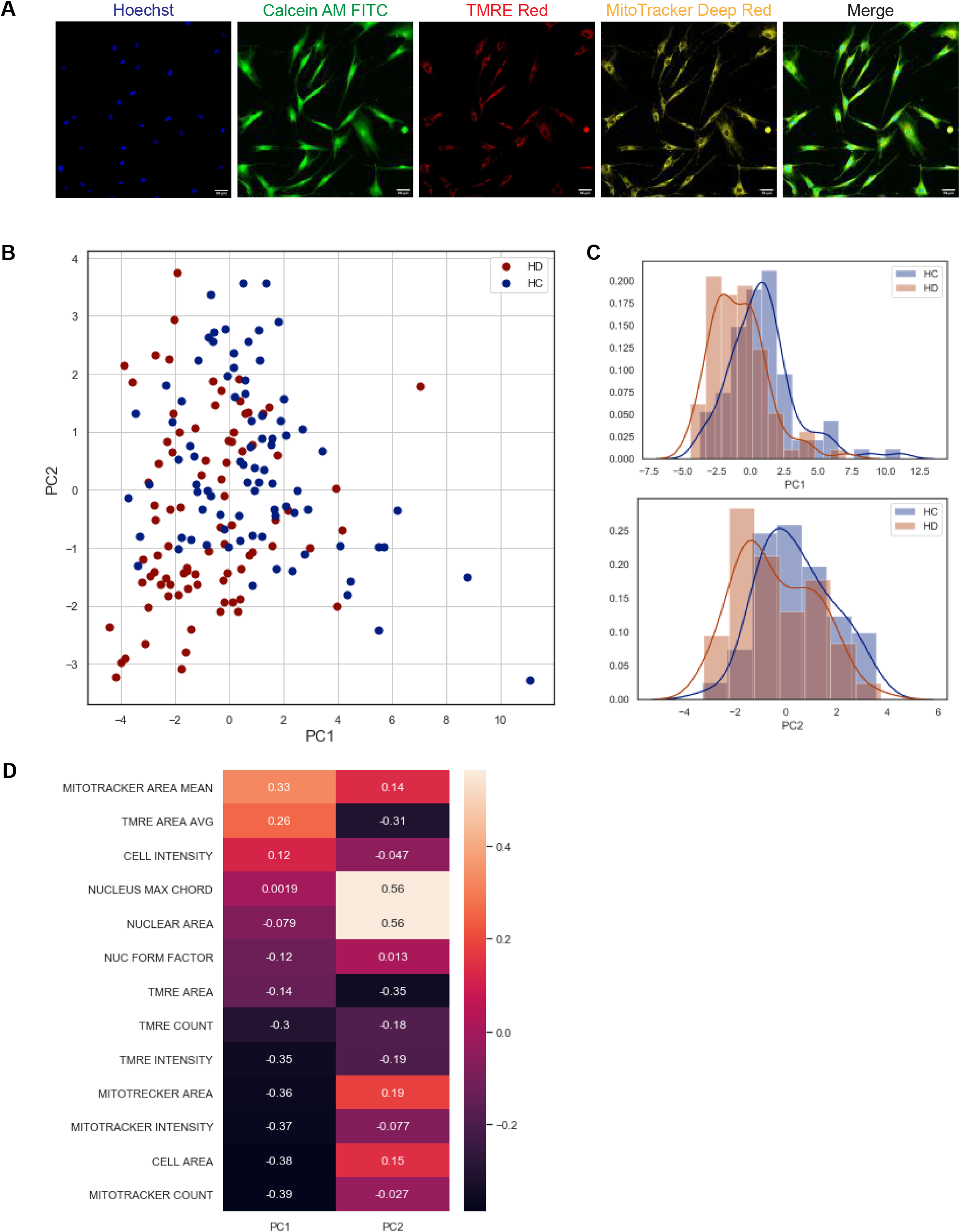
High content image based analysis of primary skin fibroblast samples of HD patients compared to HC. (A) Representative images of skin fibroblast cells stained with 4 different fluorescent vital dyes (Hoechst 22342 to label the cell nuclei, Calcein AM to label cytoplasm of living cells, TMRE to label functional mitochondria, and Mitotracker far-red to label mitochondria). The images were acquired by 20x magnification using INCell 2200 image analyzer. Scale bar = 50 µm. (B) Principal component analysis of 14HD vs. 14HC phenotypic data. (PCA graph on the right) The analysis shows a separation between the two groups. Each sample is represented by 6 circles each circle is the median of phenotypic data extracted from 20 fields coming from one well, each field contain 10-40 cells, blue circles represent HC samples, red circles represent HD samples. (Histogram graphs on the right) represent the distribution of the analyzed PCA data of the two populations in the two dimensions (PC1, PC2), blue histogram represents HC and red histogram represents HD. (D) Plot of feature importance in the two PCA dimensions PC1 & PC2, organized from the most important (orange) to the least important feature (black) to the significant difference between the two populations in the PCA analysis.

### Lack of actin cap in skin fibroblast samples from HD patients is correlated with deficient cell motility

To investigate this phenotype further, we analyzed all the nuclei images obtained from the above experiment together with newly and similarly acquired nuclear images of primary fibroblasts from five HGPS patients and respective matching controls. The HGPS cells were used as a known reference for disease nuclear morphology^23^. Results from these experiments (Figure 2A and 2B) show significant qualitative and quantitative differences in nuclear morphological features (form factor and major axis of the nucleus) between HD and HGPS and their respective adult and young HC groups. Compared to controls, the HD and HGPS cells show significantly higher roundness, which is expressed here as median nucleus form factor. In contrast, the healthy controls show a significant difference in median nucleus major axis, indicating a more oval nuclear morphology than both HD and HGPS cells.

**Fig.2.**
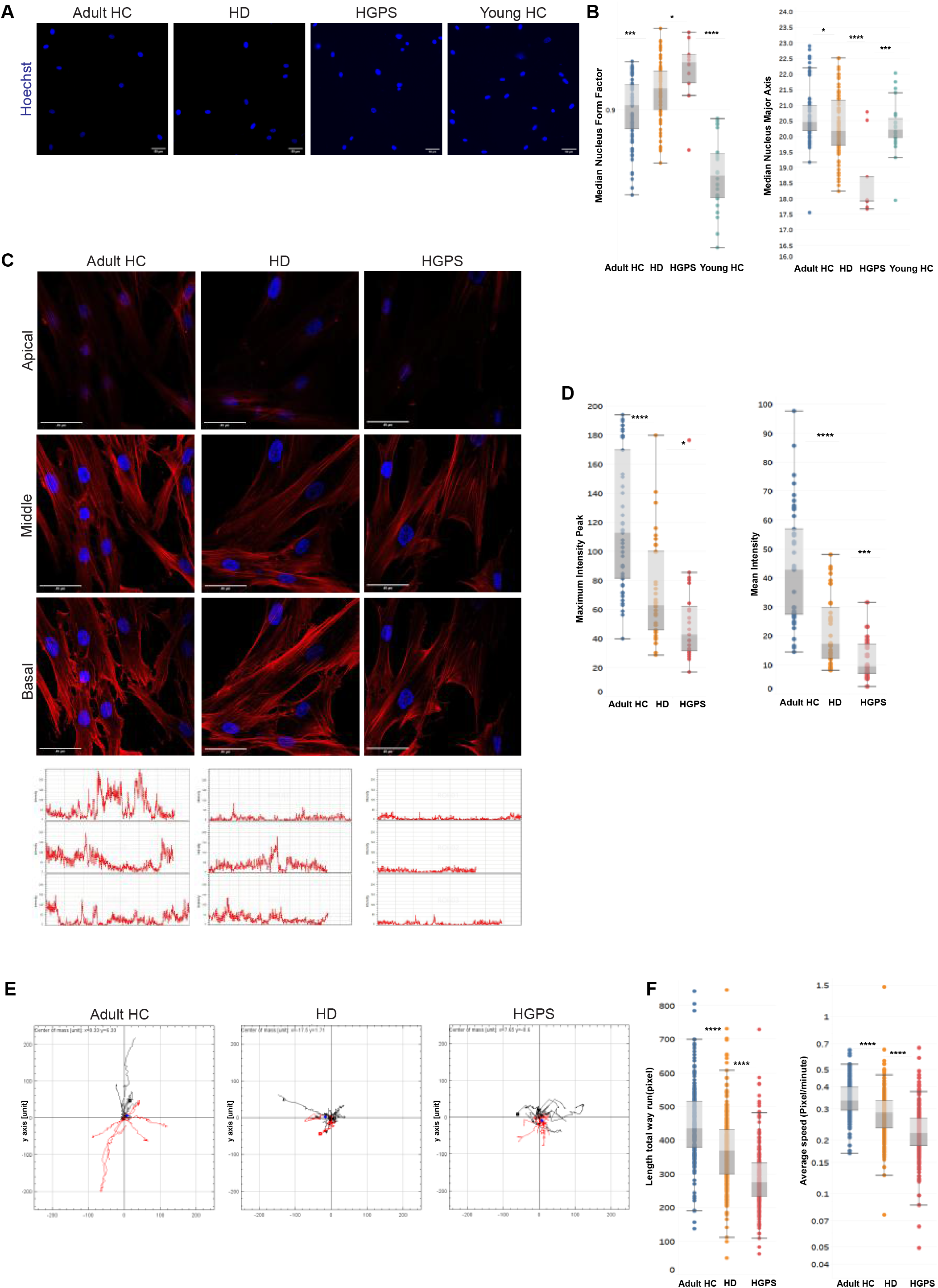
Actin Cap in HD skin fibroblast cells. (A) Representative images of fixed primary skin fibroblast cells of Adult HC, HD, HGPS, and young HC cells stained with Hoechst to label the cell nuclei and phalloidin to label F-actin, acquired with a 20x magnification using INCell 2200 image analyzer. Scale bar = 50 µm. Cells of indicated groups were segmented automatically using the INCell Developer toolbox software. (B) Mann-Whitney’s U test was performed on the medians of the nuclear form factor and nuclear major axis per well (14 HC, 14 HD, 5 HGPS, 4 young HC), ∼500 cells from each skin fibroblast sample using GraphPad 8; the results are presented in box plots using Tableau 2020.3. (C) On the upper panel representative confocal immunofluorescence staining images of HD, HC, and HGPS skin fibroblast cells of 3 z-stacks (Basal, Middle, Apical) showing less actin filaments in the apical surface of the HD and the HGPS fibroblast cells. Skin fibroblast cell images were acquired with a 40x magnification using Leica SP8 confocal microscope. Scale bar = 50μm. Cells were stained with Hoechst to label the cell nuclei and phalloidin to label F-actin of fixed cells. On the lower panel representative intensity graphs of actin in the apical z-stack, each graph represents one nucleus, Y-axis represents intensity levels (0-250), X-axis represents pixels. (D) Mann-Whitney’s U test was performed on the medians of the mean intensity of actin fibers, and of the maximum peak of different actin intensity graphs in the apical z-stack of ∼20 cells from each skin fibroblast sample (2 HC, 2 HD, 2 HGPS) showing significant lower measurements in the two parameters in HD and HGPS samples. (E) Single-cell random migration analysis of HD and HC skin fibroblast cells shows significant lower motility parameters in HD and HGPS cells compared to HC. Cell tracking analysis was carried out for 21 hours at a rate of 1 frame per 10 minutes. Several parameters of random motility were quantified on (N=2) of each HD and HC and HGPS skin fibroblast cells. Time-lapse images were acquired in brightfield with a 4x magnification using INCell 2200 image analyzer. Cells of indicated groups were tracked semi-automatically by CellTracker analysis software. After cell tracking, the cells’ paths were plotted and analyzed through The Chemotaxis and Migration Tool (ImageJ plug-in) to create the sun plots. Sun plots show the cell path for representative 15 cells from each population. (F) Mann-Whitney’s U test was performed on the medians of different parameters of random motility in HC fibroblast cells compared to the HD and HGPS per cell (2HC, 2HD, 2HGPS), ∼150 cells from each skin fibroblast sample using GraphPad 8, the results are presented in box plots using Tableau 2020.3.

Previous studies showed that nuclear morphological changes are linked to changes in the assembly or absence of a nuclear actin cap^16^. To investigate the possibility that the different nuclear morphology in HD cells is linked to an actin cap deficiency, we performed similar experiments as described above but stained the cells after 4% PFA/PBS fixation with Hoechst and phalloidin to label the nuclei and actin filaments respectively. Confocal imaging was applied to measure the phalloidin-labelled actin filaments within the stained nuclear area at three different optical z-section levels – basal, middle, and apical, taken from HD (n=2), matching HC (n=2), and HGPS (n=2) (Figure 2C). Representative images of these cells at the three different z confocal planes, along with linescan analyses of phalloidin fluorescence intensity at the apical level of the cells for three arbitrary nuclei for each group, are shown in Figure 2C upper and lower panels, respectively. Quantitative analysis of the phalloidin mean fluorescence intensity in all apical z-sections analyzed (∼50 nuclei per sample) in each group show significantly higher values in HC compared to HD and HGPS (Figure 2D). These results strongly indicate that actin filaments at the apical side of the nucleus (actin cap fibers) are more readily assembled in HC compared to HD and HGPS cells, although HD cells displayed higher levels than HGPS cells. The significantly low F-actin levels in HGPS cells are in line with their known severe actin cap deficiency^16,23^.

Based on these findings, we next tested whether the actin cap had an effect on cell motility, as shown in previous studies^17^. To this end, we performed cell motility analyses on HD (n=3), HC (n=3) and HGPS (n=3) fibroblasts samples using phase contrast time lapse microscopy and image acquisition every 10 minutes for 21 hours (tracking of ∼150 cells per sample was performed). As shown by the sunplot cell migration graphs in Figure 2E, both HD and HGPS cells display significantly lower migratory capacity compared to HC cells. Measurements of track distance and average speed of all tracked cells (Figure 2F) confirm the observed migration phenotypes and the differences between HD and HGPS as compared to HC. Altogether, this confirms that aberrant cell motility of HD cells is correlated with the newly described actin cap deficiency in these cells.

### Classification of different actin cap morphologies in HD skin fibroblast population

To further investigate the actin cap phenotype in HD cells and better classify it compared to HGPS cells, which are known for their actin cap deficiency, we developed a machine learning tool for actin cap segmentation using the Trainable Weka segmentation plugin for FIJI^24^. To this end we performed similar experiments as shown above (see Figure 2C) but images were acquired using higher resolution confocal imaging (63x 1.4NA objective and Airyscan processing) to obtain better description and higher number of the actin cap morphological features (Figure 3A). Phalloidin-labeled actin filaments within the stained nuclear area were analyzed at the apical and basal optical z-section levels from images of 120 cells in total taken from HD (n=9), matching HC (n=6), and HGPS (n=2) samples. Image analysis of HGPS cells produced no segmentation of the actin cap, as expected (Figure S3), and therefore were not included for further analyses. The data obtained from the image segmentation analysis from HC and HD cells in these experiments was analyzed using the script available at https://github.com/disc04/simplydrug. To get a better resolution in actin cap morphology in HD cells, we extracted different morphological features of the actin cap and the nucleus from the images. This enabled us to obtain classification within each group, as shown in Figure 3. PCA analysis of these features gave a 55% variance ratio showing a significant difference between the two populations (Figure 3B). The feature histogram distribution by PC dimension indicates a clear difference between the groups (Figure 3C). K-means clustering analysis of three clusters was performed, and results are shown as a heat map (Figure 3D). Cluster 0, which is composed of 23.6 % HD cells and 76.4% HC cells, is identified by moderate size and oval nuclei, with the highest actin cap area and highly parallel actin cap fibers with high F-actin intensity levels. Cluster 1, which is composed of 89.3% HD cells and 10.7% of HC cells, is characterized by the largest, more circular nuclei, together with the smallest actin cap area in combination with highly parallel actin cap fibers and low F-actin intensity levels. Cluster 2, which is composed of 70.3% HD cells and 29.7% HC cells, is characterized by the smallest and most circular nuclei, together with a larger actin cap area, compared to those in cluster 1, with non-parallel actin cap fibers and moderate F-actin intensity levels. Overall, these results show that the HD cell population is represented in two different clusters, strongly indicating that the HD actin cap phenotype is heterogeneous, although most of the HD cells analyzed were found in cluster 2. To better understand the different HD actin cap clusters, we analyzed six features describing the actin fiber network at the apical side of the nucleus. These analyses show high correlation between the features (Figure S4A,B) which enabled us to reduce the features into one representative feature of the actin cap, namely, actin network complexity. As shown in Figure S4C, this feature manages to segregate HD from HC groups, by medium and high values in 11.4% of the HD data and 15.9% of the HC data, respectively. Moreover, low values of actin network complexity are represented in 88.6% of the HD data and 84.1% of the HC data, respectively. These results suggest that the actin network complexity of HD cell population is lower than that of the HC group.

**Figure 3:**
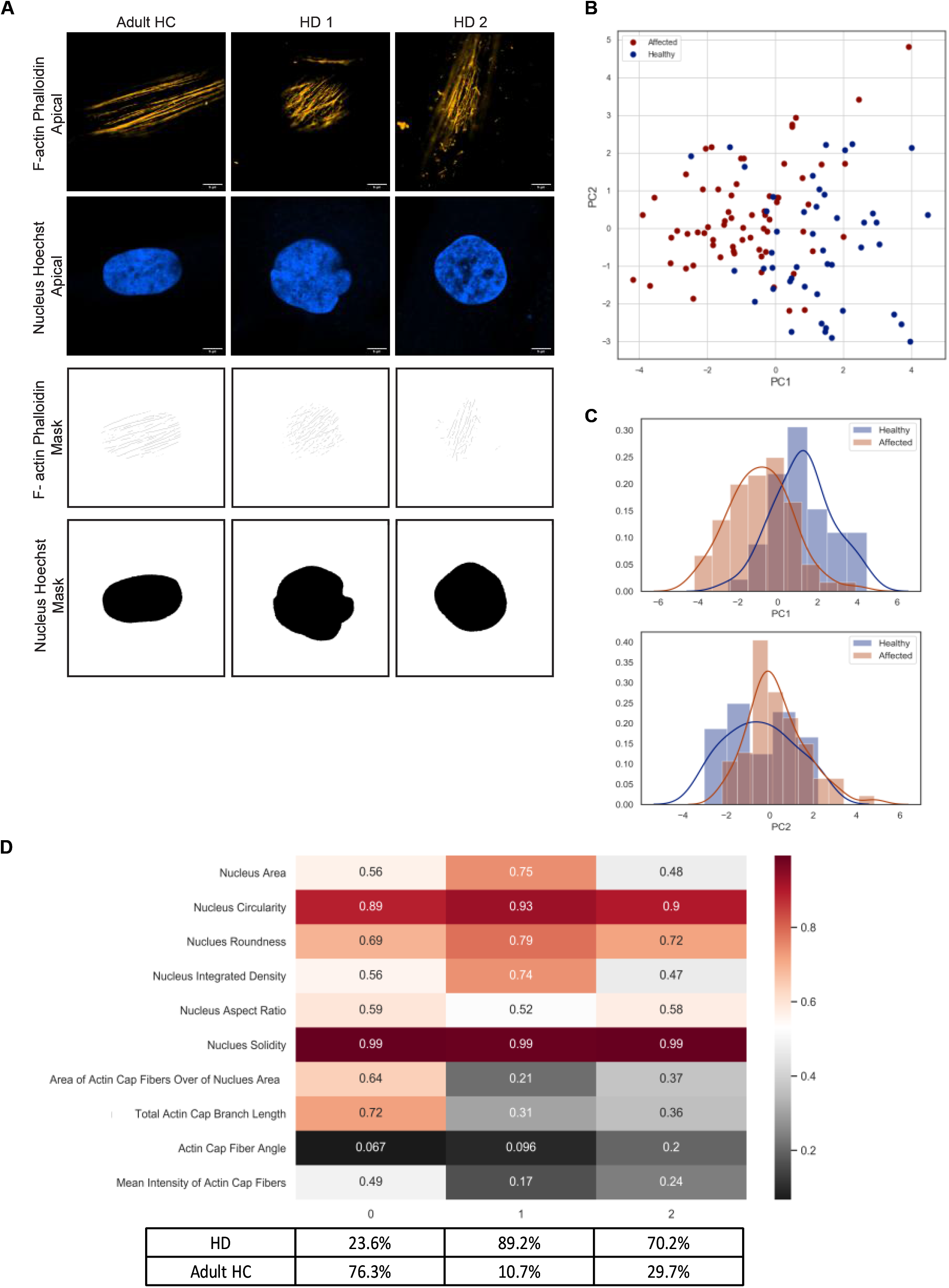
Deeper analysis for the different actin cap morphologies in HD skin fibroblast cells. (A) Representative confocal immunofluorescence staining images of data acquired under 63X magnification 1.4NA objective and Airyscan module. Cells were stained with Hoechst to label the cell nuclei and phalloidin to label actin filaments of fixed cells. Scale bar = 5μm. for analysis of different morphological clusters of actin cap, for each skin fibroblast cell 3 images were acquired – basal nucleus, apical nucleus, and apical actin. Representative Images of the segmented actin cap fibers and nucleus generated by the WEKA trainable segmentation tool. (B) PCA analysis of 8 HD vs. 8 HC actin cap and nucleus morphological features. The analysis shows a significant separation between the two groups. Each sample is represented by 10 morphological features extracted from one cell, in total ∼50 cells from each population, blue circles represent HC samples, red circles represent HD samples. (C) Histogram graphs represent the distribution of the analyzed PCA data of the two populations in the two dimensions (PC1, PC2), blue histogram represents HC and red histogram represents HD. (D) K-means analysis of three clusters of the same data analyzed by PCA in (B). The values in the plot are the average of the different values of one measurement in specific cluster Cluster 0 accumulated mainly by HC (Class0: 76.3% HC, 23.6% HD) and Cluster 1, and 2 accumulated mainly by HD cells (Cluster 1: 89.2% HD, 10.7% HC; Cluster 2: 70.2% HD, 29.7% HC).

Altogether, these results strongly support the notion that actin cap deficiency in HD is a novel representative cellular phenotype of HD primary fibroblasts, which is characterized differently within the HD sample population as compared to control.

### Image-based high content analysis algorithm for extraction of actin cap like features from 2D images

The above results demonstrate a characteristic actin cap deficiency in HD cells. We therefore wished to test whether this phenotype is linked to global actin cell organization and nuclear morphology in the cells, which could be used as a high throughput drug screening assay for HD. To this end we developed a 2D image analysis tool to extract separate features that describe the nuclear morphology and the total actin fiber organization in the cells. Since nuclear elongation and actin cap organization were highly linked (e.g., cluster 0 in Figure 3D), we focused on nuclear circularity and the standard deviation of actin fibers slopes (see the schematic representation of the image analysis tool in Figure 4A; the image analysis tool is available at https://gitlab.com/maydanw/CellDoctor). This method allowed us to increase the number of cells tested per sample, permitting HCA of these two linked features in the cells. The experiments were performed as described above (see Figure 2) and image analysis of ∼2000 cells per sample of 14 HD, 10 HC, 6 HGPS and 3 young HC fibroblasts was performed. Box plot results of the image analysis data representing each of these features is shown in Figure 4B. Using this tool, HD cells show significantly higher values of nuclear circularity compared to matching HC, which again is consistent with the levels of HGPS cells that are higher as compared to their matching HC (see Figure 2B). Moreover, HD cells show higher standard deviation of actin fiber slopes which represent a more disorganized actin fiber arrangement in the cells as compared to matching HC, in accordance with what is shown in Figure 3D regarding clustering of the actin cap content and nuclei morphology. Together with this, HGPS cells show higher organization of the actin fibers in the cells, confirming the above detailed confocal data analysis (see cluster 1 in Figure 3D), which shows that the fewer the number of actin cap fibers, the more organized they are. Therefore, in a similar way here in Figure 4B, the standard deviation of the actin fiber slopes is lower in the matching HC compared to HD and HGPS. Altogether, these results validate the use of our image HCA tool for actin content and organization in primary skin fibroblasts of HD patients to identify actin cap related phenotype in these cells.

**Figure 4:**
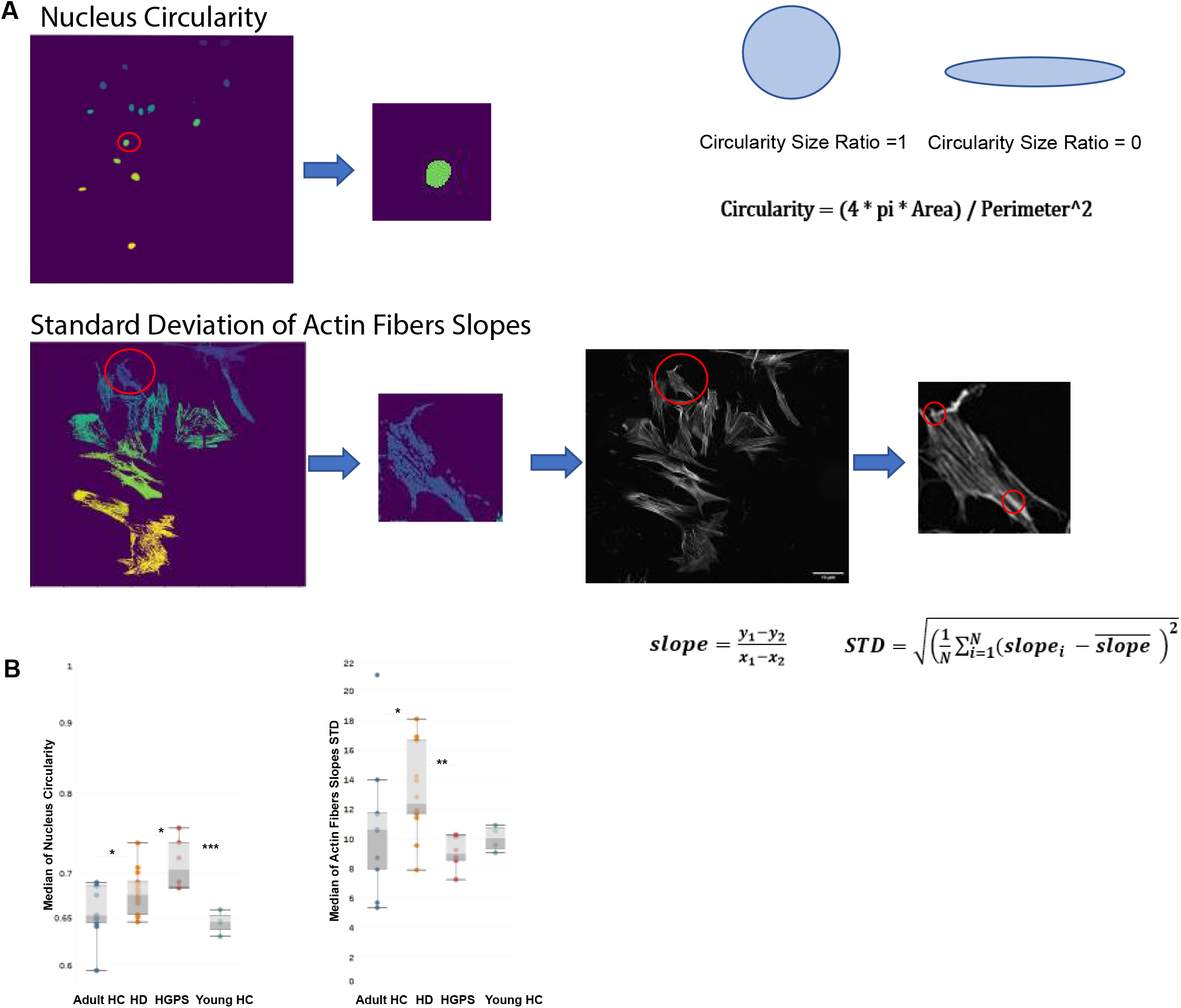
An automatic image based high content analysis algorithm for actin cap like features. (A) Schematic Representation of the segmentation steps for each feature. For the “Nucleus Circularity” feature the algorithm counts the total number of pixels constructing each nucleus segment over the nucleus image. A binary mask of each nuclear segment is generated and then used to measure the circularity. For the “Standard Deviation of Actin Fibers Slopes” feature, the algorithm segments the cell body (cytoplasm), and a binary mask is applied over the actin fibers image to receive an output image of actin fibers that overlap the specific cell’s cytoplasm. Then, an algorithm for line detection was applied on the output image in order to detect edge points couples of each actin fiber over the cytoplasm. For each actin fiber (“line”) over the cytoplasm, a linear slope was calculated, and a standard deviation value of all slopes was determined to assess the level of actin fibers parallelization over the cytoplasm. (B) Mann-Whitney’s U test was performed on the medians of different actin cap like features generated from the image analysis tool mentioned above in HC fibroblast cells compared to the HD and HGPS per patient (10 HC, 14 HD, 5 HGPS, 3 young HC), ∼500 cells from each skin fibroblast sample using GraphPad 8, the results are presented in box plots using Tableau 2020.3.

These results prompted us to test the robustness of the distinctive actin cap HD phenotype as a new predictive tool by treating the cells with two different pharmacological agents, latrunculin B that blocks actin cap formation at low concentrations^17^, and Mitoquinone mesylate (MitoQ), an antioxidant agent targeted to mitochondria to protect from oxidative stress. For functional treatment effects we performed in parallel cell motility time lapse microscopy assays as described (see Figure 2E). We hypothesized that latrunculin B treatment of HD cells would lead to a phenotype similar to that of HGPS cells that lack an actin cap completely. In contrast, we hypothesized that MitoQ treatment might recover the actin cap phenotype in HD cells due to the potential improvement in the known mitochondrial distress in HD cells^25^. To this end we performed experiments similar to those described above, where HD cells from 5 patients were treated for 24 h with either 60 nM latrunculin B or with 1 µM MitoQ, or DMSO (1:1000 v/v) as control. Matching HC (n=3) and HGPS (n=2) fibroblasts were used as controls. The effects of latrunculin B and MitoQ treatments on actin and nuclei morphology features in HD fibroblast cells are shown in Figure 5A. Interestingly, these two different drugs exert an opposite effect on the cells, where Latrunculin B treatment resembles HGPS nuclear circularity and shows a significant difference from HD DMSO control cells. In contrast, MitoQ led to a significant effect on nuclei circularity in HD treated cells compared to DMSO and similar to those of HC cells. In addition, Latrunculin B decreased the actin fiber slope standard deviation of HD cells compared to the DMSO control, indicating more parallel actin fiber organization despite the decrease in intensity. This effect resembles the phenotype that defines cluster 1 in Figure 3D. MitoQ in these experiments also affected the actin fiber slope standard deviation of HD cells significantly, but in contrast the phenotype resembles cluster 0 in Figure 3D which is representative of the HC.

**Figure 5:**
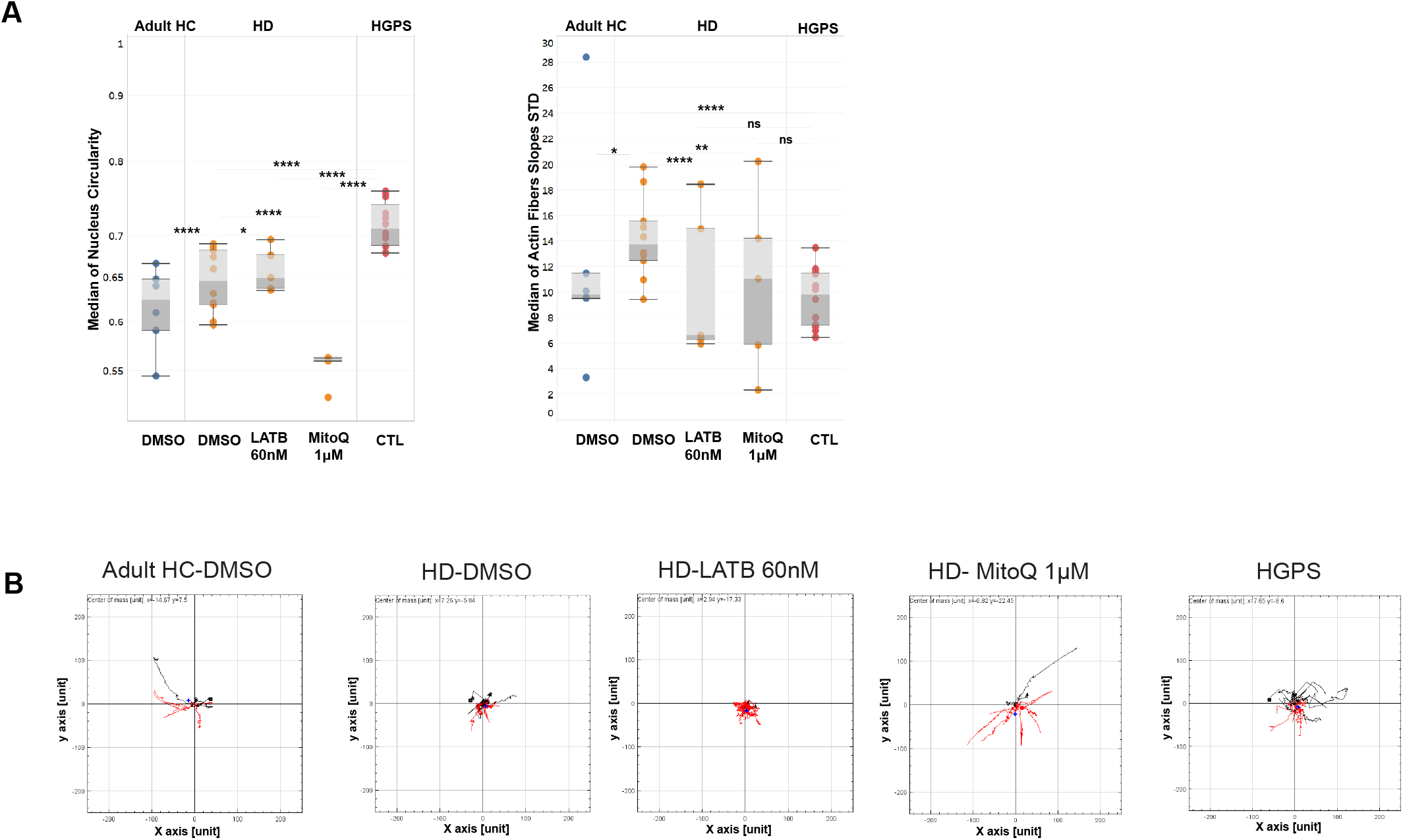
Effect of MitoQ and Latrunculin B on the actin cap in HD Fibroblast compared to HC and HGPS. (A) Mann-Whitney’s U test was performed on the medians of different actin cap like features generated from the image analysis tool mentioned above in HC fibroblast cells compared to the HD and HGPS per field (3 HC, 5 HD, 2 HGPS), ∼200 cells from each skin fibroblast sample using GraphPad 8, the results are presented in box plots using Tableau 2020.3. (B) Cell tracking analysis was carried out in the same conditions mentioned in materials and methods section. Sun plots show the cell path for representative ∼10 cells from each population. Graphs were generated by The Chemotaxis and Migration Tool (ImageJ plug-in).

Altogether, from these results we conclude that Latrunculin B at low concentrations exacerbates the actin cap phenotype of the HD cells, making it more similar to the actin cap phenotype of HGPS cells. In contrast, MitoQ improves the actin cap phenotype significantly in HD cells, making it more similar to the actin cap phenotype of HC cells. Remarkably, MitoQ also restored the motility of HD cells in all the measured parameters whereas Latrunculin B did not (see Figure 5B). Moreover, the opposite effects of the two drugs on cell motility demonstrate the direct functional link between their respective effects on actin, actin cap, and nuclear morphologies and cell motility. Altogether, these results strongly support the use of the actin cap associated features and cell motility assay to screen and evaluate drugs on HD patient cells for potential personalized treatment of HD patients in the clinic.

## Discussion

Although the HD mutation in the Htt gene is a well-characterized cause of disease, the mechanism of disease in the different tissues and organs remains to be elucidated. Brain pathology has become a hallmark of HD, but critical mass of new studies suggest that HD is a multi-system challenging disorder, contributing to the patient’s individual disease progression and disease severity independently of the high number of CAG repeats in the mtHtt^7,8^. Therefore, it is important to characterize the phenotypic implications of mtHtt expression in cells other than neurons, such as primary HD fibroblasts, and try to connect them with specific cellular biological functions affected in the HD patients’ cells. To this end we applied an unbiased high throughput cell-based image analysis approach to characterize an HD phenotype in these cells. Our results show that primary HD patients’ fibroblasts exhibit a distinctive cellular phenotype that allows accurate classification from HC by image-based HCA tools. This analysis on 16 HD fibroblasts samples compared to 18 matching HC, strongly supports the robustness of such phenotypic classification. The HD cell phenotype is characterized by differences in cell morphology and mitochondrial features previously described in other HD models^25,26^. In addition, we found, for the first time, that significant nuclear morphological features are distinctive in the HD cell population (see Figure 1). A deeper analysis into the nuclear morphology of the cell by confocal microscopy allowed us to test the known link between nuclear roundness and the actin cap morphology (Figure 2)^16^. These experiments were performed using both HC fibroblasts as positive control and HGPS fibroblasts as reference for actin cap depletion control, since these patient cells are known to have a severe actin cap and concomitant nuclear morphology abnormalities^20^. The confocal morphometric results strongly indicate that HD cells resemble more the HGPS nuclear morphology together with an associated actin cap deficient phenotype than the HC, suggesting that HD shares aspects associated with HGPS laminopathy. Further support for the HD phenotype comes from our mechanistic actin cap-related experiments, measuring cell migration for 24 hours by time-lapse microscopy (Figure 2)^17^. The migratory capacity of the HD cells confirms that their actin cap deficiency is correlated with their cell motility, which falls also in between the HC and HGPS control groups.

Studies show that mutant Htt interferes with actin dependent cellular remodeling. Mutant Htt fragments interact abnormally with actin binding proteins in HEK293 cells transfected with Htt Exon 1^27^, and in mouse striatal neuron-derived cell line (STHdh) expressing full-length endogenous levels of either wild-type (STHdhQ7/Q7), or mutant (STHdhQ111/Q111)^28^. Different studies support the idea that Htt is involved in regulating adhesion and actin dependent functions including those involving α-actinin^29^ – an actin-binding protein and one of the components of the actin cap^30,31^. α-actinin is recruited to focal complexes, where it provides a structural link between integrins and actin microfilaments^31,32^. An interactome analysis of huntingtin in mouse brain models showed that Htt is associated with α-actinin-1, -2 and -4 both in wild type and Htt mutant huntingtin^33,34^. Follow up studies validated these results through immunoprecipitations using exogenously expressed proteins,immunofluorescence co-localization, and proximity ligation assay, showing a functional interaction of Huntingtin with isoforms of α-actinin in human primary fibroblasts and neurons^29^. This study, along with work of others^35,36,37^, proposed a model in which Huntingtin regulates α-actinin-1 localization and couples growth factor signaling with actin polymerization with bundling functions at new sites of adhesion. This evidence regarding htt and α-actinin subtypes strongly supports the possibility in HD patients’ fibroblasts that actin cap might be aberrated due to abnormal binding between α-actinins and htt due to mt-htt expression.

Lamin B protein is one of the two subgroups that compose the lamin family which not only provide structural support to the nuclear envelope membrane, but is also involved in a wide variety of cell functions and processes^38^. Lamin B is expressed in almost all cell types independently of their differentiation state^39^, suggesting its critical role in mammalian cell survival^40^. It is also known that lamin B binds directly to F-actin that forms the actin cap^41^. Thus, altered levels of lamin B may have a direct effect on the formation, stabilization, and organization of the actin cap. Very recent work demonstrated a new pathogenic mechanism for HD by showing an increase in the Lamin B1 protein levels in HD brains in a neuron-type specific manner, which is correlated with alterations in nuclear morphology and nucleocytoplasmic transport^42^.

We also show here that the actin cap phenotype in HD fibroblast cells is correlated with a distinctive nuclear morphology which allowed us to develop an image high throughput analysis tool of the actin content and organization in the nucleus area and nuclear morphology phenotype using 20X low power magnification. This analysis allowed us to characterize the HD cell populations described in distinctive clusters that are representative of the actin cap phenotype status, showing a more severe phenotype (like HGPS cells) with the roundest nuclei and complete actin cap depletion with a parallel organization of the actin cap fibers (cluster 1), while other HD cells have a round nucleus with disorganized actin cap instead (cluster 2). The majority of the analyzed HD cells were clustered under cluster 1 (mild phenotype) while most of the HC cells integrated cluster 2 (normal actin cap with oval like nuclei phenotype). The relative simplicity of this method should allow using this HD cell phenotype biomarker as reliable indicator to test or screen drug effects for patients with HD. This observation could allow us to classify/treat the patients in the future according to their cell phenotype severity level. The use of known pharmacological agents such as Latrunculin B and MitoQ to test their expected opposite effects on the HD actin content and organization in the nucleus area and nuclear morphology allowed us to test the drug screening potential of the assay. From these experiments, it becomes clear that while Latrunculin B exacerbates the HD cell phenotype by blocking actin polymerization, MitoQ, which reduces oxidative stress, improves significantly all the morphological features as well as cell migration parameters in the HD cells. Interestingly, these results strongly indicate that the mitochondrial function may regulate the actin cap/nuclear system and concomitant cell motility. Aberrant mitochondrial activity, possibly by increased reactive oxygen species (ROS) production, may be responsible for the actin cap phenotype in HD cells which can be reversed by MitoQ treatment that also recovers functional motility in HD patients’ cells.

Our results strongly support the power of the image based phenotypic HCA assay in discovering new biomarkers in HD. In addition, they also support the use of primary skin fibroblasts from HD patients as personalized model for disease phenotype classification based on linked actin cap/nuclear morphologies that functionally affect cell migration, which can be recovered by pharmacological agents. Altogether, our results and the HCA phenotypic tool used to describe the HD cell phenotype open the door for future drug testing or drug screening campaigns directly using the HD patient derived cells.

## Materials and Methods

Most of the reagents used in this study (otherwise stated) were purchased from Thermo Fischer Scientific, USA.

### Primary skin fibroblasts samples and cell culture handling

Skin fibroblasts (see Table 1) were isolated from 28 individuals (age 38-68), 14 HD patients and 14 matched healthy controls, the HD skin fibroblast samples were purchased from Coriell Institute in New Jersey and the healthy controls samples were sent from the neurology department at the University of Michigan medical center under signed Material transfer agreement. Samples were then expanded in culture media DMEM supplemented with 1% MEM Sodium Pyruvate, 1% PSA (Biological industries, Israel), 10% heat inactivated FBS, and 1% NEAA in polystyrene plastic 75-cm2 culture flasks (Corning, NY) at 37°C with 5% CO2. Cells passages and expansion of human skin fibroblasts were performed when cells were at 80-100% confluency. Cells passaging and re-plating was accomplished using 0.25% Trypsin-EDTA (Biological industries, Israel) for 2 minutes, followed by addition of twice the volume of a complete culture media to neutralize the enzyme. Cells were subsequently centrifuged at 1200 rpm for 5 minutes before their pellets being resuspended in 1 ml medium for cell counting using TC10 automated cell counter (BioRad USA) before re-plating. All experiments were performed between passages 7– 15.

**Table 1:** List of skin fibroblasts samples age and gender matched from HD and HC individuals

| HD individual | Gender | Age | HC individual | Gender | Age |
| --- | --- | --- | --- | --- | --- |
| GM06274 | Female | 56 YR | 0579C | Male | 40 YR |
| GM04196 | Female | 51 YR | 0233C | Female | 61 YR |
| GM04200 | Male | 53 YR | 0848C | Female | 48 YR |
| GM04212 | Female | 50 YR | 1170C | Male | 43 YR |
| GM04287 | Male | 43 YR | 0561C | Male | 48 YR |
| GM04476 | Male | 57 YR | 0795C | Male | 52 YR |
| GM04709 | Female | 38 YR | 0553C | Male | 49 YR |
| GM04715 | Male | 40 YR | 0769C | Male | 46 YR |
| GM04717 | Female | 44 YR | 0143C | Male | 42 YR |
| GM04719 | Female | 39 YR | 0351C | Male | 62 YR |
| GM04721 | Female | 37 YR | 0981C | Male | 50 YR |
| GM04799 | Male | 47 YR | 0536C | Male | 68 YR |
| GM04819 | Male | 48 YR | 0205C | Female | 68 YR |
| GM04887 | Female | 48 YR | 0025C | Male | 48 YR |

### Image based cell HCA live phenotyping experiments

For live image microscopy experiments, 1500 cells in 100μl of complete medium per well were automatically plated in 96-well plates (Grenier, Austria) using Tecan Freedom EVO 200 robot equipped with a 96 MultiChannel Arm (MCA). The plates accommodated serially and in columns, alternating 5 cell samples from each group HD and HC. All steps of the seeding, washing, staining, replacing media protocols were programmed for running automatically with optimized pipetting parameters for each task. After 24 hours incubation at 37 °C and 5 % CO2, culture media was removed and replaced, after one wash with DPBS, with fluorescent dyes mix diluted in HBSS. After 30 minutes incubation at 37°C, and 5% CO2 the plate was transferred to the IN Cell Analyzer 2200 (GE Healthcare) for image acquisition under cell culture environmental conditions. The fluorescent vital dyes mix used contain Hoechst 33342 (Merck-Sigma, USA) 1:10000, CellTrace™ Calcein Green AM (Invitrogen, USA) 1:5000, mitochondria TMRE red (Invitrogen, USA) 1:2500 and MitoTracker™ Deep Red (Invitrogen, USA) 1:5000. Twenty images per fluorescent channel (fixed spacing fields) for each well were acquired in four different channels in less than 60 min. All images under a 20x magnification were taken using the same acquisition protocol with constant exposures for each fluorescent channel for each of the four fluorescent dyes used to stain the cells. Image acquisition has been done in a horizontal serpentine pattern. The multiple cell images were subsequently segmented and high content analyzed using IN Cell Developer software (GE Healthcare).

### Actin F and nuclear staining assay for fluorescent microscopy

Cells from HC and HD samples were seeded and culture in 96 well plates as described above, then fixed in 4 % v/v Paraformaldehyde (Electron Microscopy Sciences, USA) in PBS for 10 min, and washed 3 times with DPBS. Cells were then permeabilized with 0.1% Triton X-100 (Merk-Sigma, USA) in PBS for 10 min at room temperature. Then Blocking solution 5% w/v BSA (Chem-Impex, USA) in TBST was added to the cells for 1 hour. After 1h. After 1 hour, the sample was washed 3 times with 0.05% Triton X-100 for 5 min. After the removal of the 5% BSA solution

The cells were washed with PBS and the nuclei and actin filaments were stained at room temperature for 1 hour in the dark with a mix of 30μl Hoechst 33342 1:10000 and Phalloidin (Merk-Sigma, USA) 1:400 in 5% BSA (in TBST) respectively. The stained cells were washed three times with DPBS and image acquisition and analysis of the plate was performed similarly as described above.

### Confocal microscopy, image and data analysis

Confocal microscopy of actin and nuclei labeled cells for actin cap validation experiments was performed under a Leica SP8 LIGHTNING (Leica Microsystems, Germany) confocal microscope with a 40x 1.2 N.A objective using 561 nm and 405 nm laser wavelengths for actin fibers and nucleus, respectively. The analysis of the actin cap fibers at the at the apical and basal focal planes was done using the LAS X software (Leica Microsystems, Germany). To determine different morphological clusters of actin cap in the HD skin fibroblast confocal microscopy analysis was performed using a Zeiss LSM800 confocal microscope with a 63x 1.4NA objective and Airyscan module. A region of interest of 40.6 µm by 40.6 µm and scan zoom of 2.5 was acquired using 561 nm and 405 nm laser wavelengths for actin fibers and nucleus, respectively. Images were processed using the Airyscan algorithm in the ZEN blue software (Zeiss). Actin cap fibers were identified using Trainable Weka segmentation plugin for FIJI^24^, and thresholding was used to define the nucleus region. Fiber orientation analysis was done using the Ridge Detection and Orientation J plugins for FIJI.

### Cell migration time lapse microscopy assay

The primary fibroblasts were plated in a 96-well plate at low density (800 cells/well) and then incubated overnight at 37°C and 5% CO2. Time lapse microscopy imaging of these plated cells was performed using an IN Cell Analyzer 2200 equipped with a long working distance 4× objective, phase contrast capability and controlled environmental chamber. The image acquisition frequency was set every 10 min for 21 h for 30 wells. The analysis of the time lapse images was performed by CellTracker program using semi-automated cell tracking function which allows a quantitative determination of cell motion parameters, including cell total travel length, net displacement and velocity. The cells paths were plotted and analyzed through The Chemotaxis and Migration Tool (ImageJ plug-in) to create the sun plots shown in the results.

### Custom image analysis tool for actin cap like morphological features

The image analysis tool was fully programmed and validated in our lab using python 3.7 programming language https://gitlab.com/maydanw/CellDoctor microscopy images of skin fibroblast cells were processed through the tool, extracting the following actin cap like parameters: The nucleus circularity.

The standard deviation of actin fibres’ slope in each cell –slopes of all actin fibers were calculated per cell followed by a calculation of the standard deviation of the slopes per cell.

**Circularity = (4 * pi * Area) / Perimeter^2**

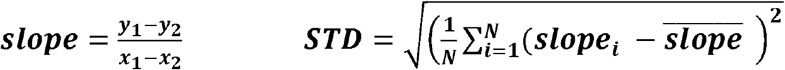

### Statistical analysis

The raw data obtained from the image analysis in this study was further statistically analyzed using the imaging assay development notebook script at https://github.com/disc04/simplydrug site, found under HTS notebook, under imaging-based assays development script being adapted to our data. for checking if the two groups were significantly different from each other, the Mann-Whitney U test was used on the medians of different parameters comparing between two groups using GraphPad 8, and the results were presented in box plots using Tableau 2020.3. for graph generations.

## Author contributions

S.G., M.W., designed the study. S.G. performed and interpreted all experiments. O.P. developed the algorithm for extraction of actin cap like features from 2D images and performed the relevant statistical analysis. L.F developed the machine learning tool for actin cap segmentation using the Trainable Weka segmentation plugin for FIJI. Maydan Weinrab developed the costumed image analysis tool found under: https://gitlab.com/maydanw/CellDoctor. A.A. developed the imaging assay development notebook script at https://github.com/disc04/simplydrug. H.W. provided guidance and help for the actin cap classification experiments. M.W. and H.W. contributed to data interpretation; and S.G., M.W. H.W. contributed to reviewing and editing the manuscript.

## Acknowledgements

This work was supported by the Taube-Koret Global Collaboration in Neurodegenerative Diseases Fund. S.G. Ph.D scholarship was partially supported by the Planning & Budgeting Committee of the Council for higher education. We acknowledge the help of Adam Baransi and Itay Bosidan from The Future Scientists Center–Alpha Program at Tel Aviv Youth University, Tel Aviv 6997801, Israel. H.W. acknowledges support from the Israel Science Foundation (1738/17) and from the Rappaport Family Foundation. H.W. is an incumbent of the David and Inez Myers Career Advancement Chair in Life Sciences.

## Supplementary Legends

<u>Figure S1: Detailed high content image-based analysis of primary skin fibroblast samples of HD patients compared to HC</u>

(A.1) Correlation heat map of morphological features of 14 HD vs. 14 HC primary skin fibroblast samples; the correlation coefficients are color-coded from deep blue (−1) to yellow (1). (B) Box plots of the average of phenotypic data per patient and per group. The data was extracted from 6 wells for each sample, each well containing 20 fields, each field containing 10-40 cells, blue represents the HC samples and red represents the HD samples. (C) Seaborn pairplot of the morphological data. The point colors correspond to the groups HD in blue and HC in orange. The plot is based on samples of 84 points coming from each group.

<u>S2: Cell migration videos</u>

(A) Representative cell migration videos with marked migration path for the analyzed cells (each cell marked with different color) of the three groups Adult HC, HD and HGPS.

<u>S3: Deeper analysis for the different actin cap morphologies in HD skin fibroblast cells</u>

Representative confocal immunofluorescence staining images of data acquired under 63X magnification 1.4NA objective and Airyscan module for HGPS primary fibroblasts in the apical plane. Cells were stained with Hoechst to label the cell nuclei and phalloidin to label actin filaments of fixed cells. Scale bar = 5μm. Actin cap fibers were not detected by WEKA segmentation tool in HGPS fibroblast.

<u>S4: Detailed analysis for actin cap morphology in HD skin fibroblast cells</u>

(A) Correlation heat map of actin cap filaments morphological features of 9 HD vs. 7 HC primary skin fibroblast samples; the correlation coefficient is color coded from deep blue (−1) to yellow (1). (B) Seaborn pairplot of the morphological data. The point colors correspond to the groups HC in blue and HD in orange. The plot is based on samples of ∼11000 points coming from each group. (C) Bar chart representing the percentage of each population HC in blue and HD in red in the branching feature divided into three classes (low, medium, high).

<u>S5: Detailed analysis for actin cap morphological cluster in HD skin fibroblast cells</u>

(A) Correlation heat map of actin cap and nucleus morphological features of 9 HD vs. 7 HC primary skin fibroblast samples; the correlation coefficient is color coded from deep blue (−1) to yellow (1). (B) Box plots of the average of phenotypic data per patient and per group. The data was extracted from ∼50 cells from each group, blue represents the HC samples and red represents the HD samples. (C) Seaborn pairplot of the morphological data. The point colors correspond to the groups HC in blue and HD in orange. The plot is based on samples of ∼50 points coming from each group. (D) Seaborn pairplot of the morphological data in 3 clusters. The point colors correspond to the clusters, cluster 0 in blue, cluster 1 in orange, and cluster 2 in green. The plot is based on samples of ∼50 points coming from each group.

## References

1. Walker, F. O. Huntington’s disease. Lancet 369, 218–228 (2007).

2. Kirkwood, S. C., Su, J. L., Conneally, P. M. & Foroud, T. Progression of symptoms in the early and middle stages of Huntington disease. Arch. Neurol. 58, 273–278 (2001).

3. Strong, T. V. et al. Widespread expression of the human and rat Huntington’s disease gene in brain and nonneural tissues. Nat. Genet. 5, 259–265 (1993).

4. Dé Ric Saudou, F. & Humbert, S. The Biology of Huntingtin. (2016). doi:10.1016/j.neuron.2016.02.003

5. Harjes, P. & Wanker, E. E. The hunt for huntingtin function: interaction partners tell many different stories. Trends Biochem. Sci. 28, 425–433 (2003).

6. Moily, N. S. et al. Transcriptional profiles for distinct aggregation states of mutant Huntingtin exon 1 protein unmask new Huntington’s disease pathways. Mol. Cell. Neurosci. 83, 103–112 (2017).

7. Sathasivam, K. et al. Formation of polyglutamine inclusions in non-CNS tissue. Hum.Mol. Genet. 8, 813–822 (1999).

8. Mielcarek, M. Huntington’s disease is a multi-system disorder. Rare Dis. 3, e1058464 (2015).

9. Connolly, G. P. Fibroblast models of neurological disorders: Fluorescence measurement studies. Trends Pharmacol. Sci. 19, 171–177 (1998).

10. Huang, H.-M. et al. Use of Cultured Fibroblasts in Elucidating the Pathophysiology and Diagnosis of Alzheimer’s Diseasea. Ann. N. Y. Acad. Sci. 747, 225–244 (2006).

11. Marchina, E. et al. Gene expression profile in fibroblasts of Huntington’s disease patients and controls. J. Neurol. Sci. 337, 42–46 (2014).

12. Gardiner, S. L. et al. Bioenergetics in fibroblasts of patients with Huntington disease are associated with age at onset. Neurol Genet 4, 275 (2018).

13. Claudia S. Alge, †, Stefanie M. Hauck, ‡, Siegfried G. Priglinger, †, Anselm Kampik,†and & Marius Ueffing*, ‡. Differential Protein Profiling of Primary versus Immortalized Human RPE Cells Identifies Expression Patterns Associated with Cytoskeletal Remodeling and Cell Survival. (2006). doi:10.1021/PR050420T

14. Wald-Altman, S., Pichinuk, E., Kakhlon, O. & Weil, M. A differential autophagy-dependent response to DNA doublestrand breaks in bone marrow mesenchymal stem cells from sporadic ALS patients. DMM Dis. Model. Mech. 10, 645–654 (2017).

15. Solmesky, L. & Weil, M. Personalized Drug Discovery: HCA Approach Optimized for Rare Diseases at Tel Aviv University. Comb. Chem. High Throughput Screen. 17, 253–255 (2014).

16. Khatau, S. B. et al. A perinuclear actin cap regulates nuclear shape. Proc. Natl. Acad. Sci. U. S. A. 106, 19017–22 (2009).

17. Kim, D. H., Cho, S. & Wirtz, D. Tight coupling between nucleus and cell migration through the perinuclear actin cap. J. Cell Sci. 127, 2528–2541 (2014).

18. Kim, D. H., Chambliss, A. B. & Wirtz, D. The multi-faceted role of the actin cap in cellular mechanosensation and mechanotransduction. Soft Matter 9, 5516–5523 (2013).

19. Zink, D., Fischer, A. H. & Nickerson, J. A. Nuclear structure in cancer cells. Nat. Rev. Cancer 4, 677–687 (2004).

20. Booth-Gauthier, E. A. et al. Hutchinson-Gilford progeria syndrome alters nuclear shape and reduces cell motility in three dimensional model substrates. Integr. Biol. (United Kingdom) 5, 569–577 (2013).

21. Kim, J. K. et al. Nuclear lamin A/C harnesses the perinuclear apical actin cables to protect nuclear morphology. Nat. Commun. 8, 1–13 (2017).

22. Costa, V. & Scorrano, L. Shaping the role of mitochondria in the pathogenesis of Huntington’s disease. EMBO Journal 31, 1853–1864 (2012).

23. Hale, C. M. et al. Dysfunctional connections between the nucleus and the actin and microtubule networks in laminopathic models. Biophys. J. 95, 5462–5475 (2008).

24. Arganda-Carreras, I. et al. Trainable Weka Segmentation: a machine learning tool for microscopy pixel classification. Bioinformatics 33, 2424–2426 (2017).

25. Carmo, C., Naia, L., Lopes, C. & Rego, A. C. Mitochondrial dysfunction in huntington’s disease. in Advances in Experimental Medicine and Biology 1049, 59–83 (Springer New York LLC, 2018).

26. Squitieri, F. et al. Abnormal morphology of peripheral cell tissues from patients with Huntington disease. doi:10.1007/s00702-009-0328-4

27. Munsie, L. et al. Mutant huntingtin causes defective actin remodeling during stress: Defining a new role for transglutaminase 2 in neurodegenerative disease. Hum. Mol. Genet. 20, 1937–1951 (2011).

28. Angeli, S., Shao, J. & Diamond, M. I. F-actin binding regions on the androgen receptor and huntingtin increase aggregation and alter aggregate characteristics. PLoS One 5, e9053 (2010).

29. Tousley, A. et al. Huntingtin associates with the actin cytoskeleton and α-actinin isoforms to influence stimulus dependent morphology changes. PLoS One 14, e0212337 (2019).

30. Djinovic’ s, K. D.-, Carugo, D.-, Young, P., Gautel, M. & Saraste, M. Structure of the-Actinin Rod: Molecular Basis for Cross-Linking of Actin Filaments-Actinin is composed of an amino-terminal actin-binding region consisting of two calponin homology (CH) domains, a central rod containing four spectrin-like re. Cell 98, (1999).

31. Sjöblom, B., Salmazo, A. & Djinović-Carugo, K. α-Actinin structure and regulation.Cellular and Molecular Life Sciences 65, 2688–2701 (2008).

32. Meacci, G. et al. alpha-Actinin links extracellular matrix rigidity-sensing contractile units with periodic cell-edge retractions. Mol. Biol. Cell 27, 3471–3479 (2016).

33. Culver, B. P. et al. Proteomic analysis of wild-type and mutant huntingtin-associated proteins in mouse brains identifies unique interactions and involvement in protein synthesis. J. Biol. Chem. 287, 21599–21614 (2012).

34. Shirasaki, D. I. et al. Network organization of the huntingtin proteomic interactome in mammalian brain. Neuron 75, 41–57 (2012).

35. Greenwood, J. A., Theibert, A. B., Prestwich, G. D. & Murphy-Ullrich, J.E. Restructuring of focal adhesion plaques by PI 3-kinase: Regulation by PtdIns (3,4,5)-P3 binding to α-actinin. J. Cell Biol. 150, 627–641 (2000).

36. Kovac, B., Teo, J. L., Mäkelä, T.P. & Vallenius, T. Assembly of non-contractile dorsal stress fibers requires α-actinin-1 and Rac1 in migrating and spreading cells. J. Cell Sci. 126, 263–273 (2013).

37. Guo, F., Debidda, M., Yang, L., Williams, D. A. & Zheng, Y. Genetic deletion of Rac1 GTPase reveals its critical role in actin stress fiber formation and focal adhesion complex assembly. J. Biol. Chem. 281, 18652–18659 (2006).

38. Hozak, P., Sasseville, A. M. J., Raymond, Y. & Cook, P. R. Lamin proteins form an internal nucleoskeleton as well as a peripheral lamina in human cells. J. Cell Sci. 108, 635–644 (1995).

39. R.M. Verstraeten V.V., Broers, J.S. Ramaekers, F. & M. van Steensel, M. The Nuclear Envelope, a Key Structure in Cellular Integrity and Gene Expression. Curr. Med. Chem. 14, 1231–1248 (2007).

40. Harborth, J., Elbashir, S. M., Bechert, K., Tuschl, T. & Weber, K. Identification of essential genes in cultured mammalian cells using small interfering RNAs. J. Cell Sci.114, 4557–4565 (2001).

41. Simon, D. N., Zastrow, M. S. & Wilson, K. L. Direct actin binding to A- and B-type lamin tails and actin filament bundling by the lamin A tail. Nucleus 1, 264–272 (2010).

42. Alcalá□Vida, R. et al. Neuron type□specific increase in lamin B1 contributes to nuclear dysfunction in Huntington’s disease. EMBO Mol. Med. 13, e12105 (2021).

