## Supplementary figures and images for "Perturbed actin cap and nuclear morphology in primary fibroblasts of Huntington’s disease patients as a new phenotypic marker for personalized drug evaluation"

### Supplementary 1

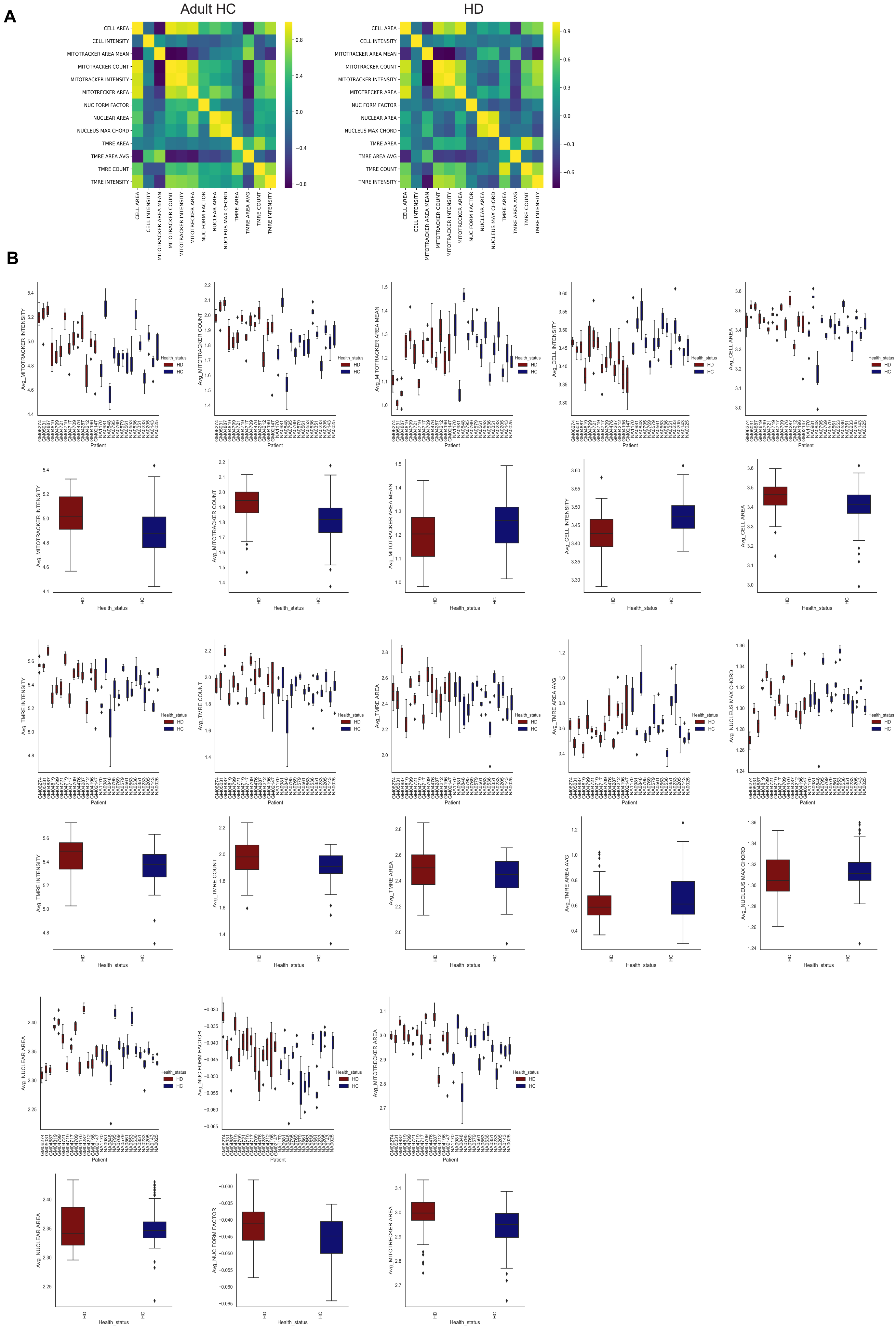

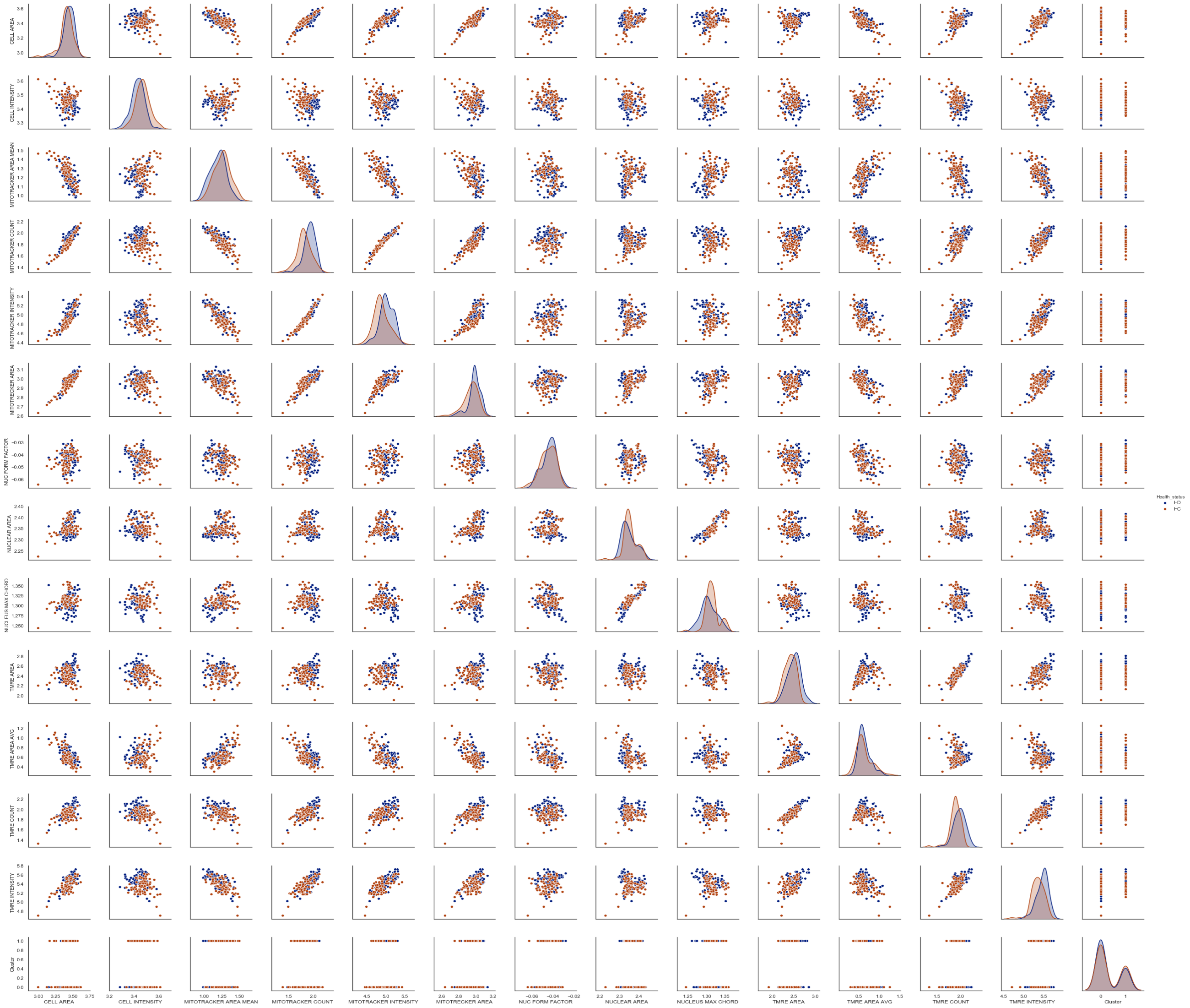

### Supplementary 2

#### Slide 1
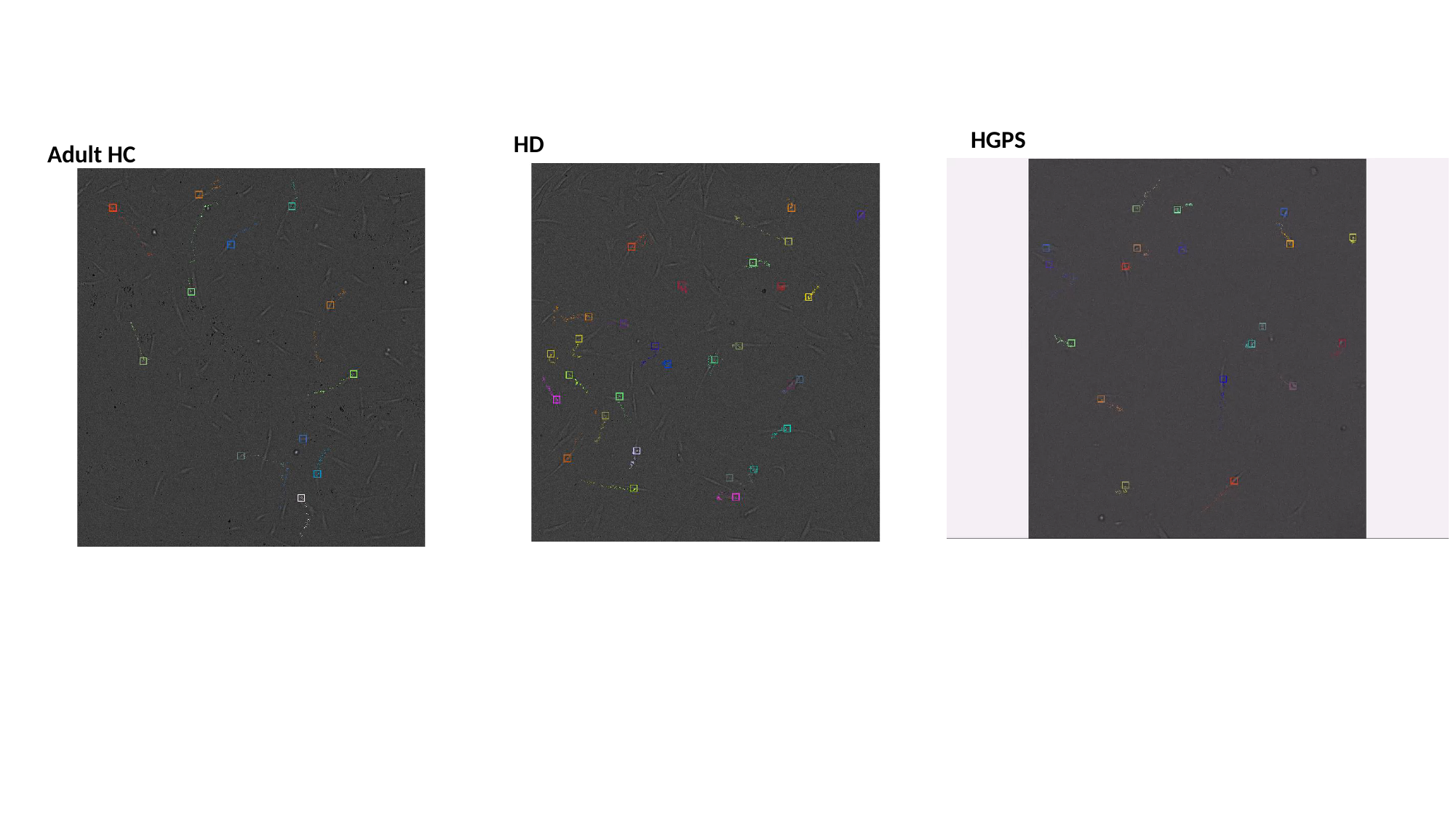

HGPS
HD
Adult HC

### Supplementary 3

A

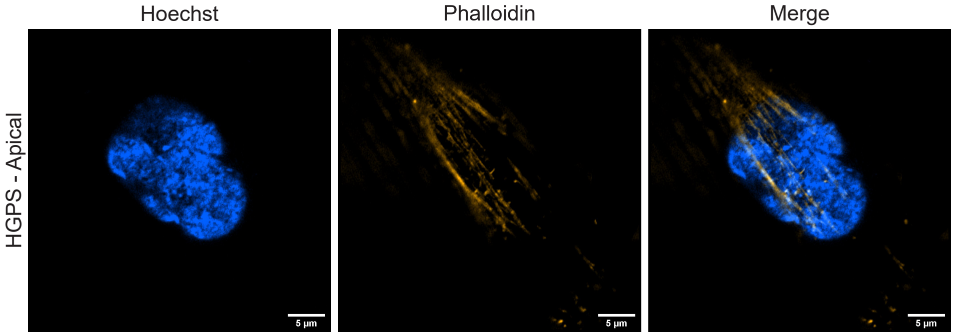

### Supplementary 4

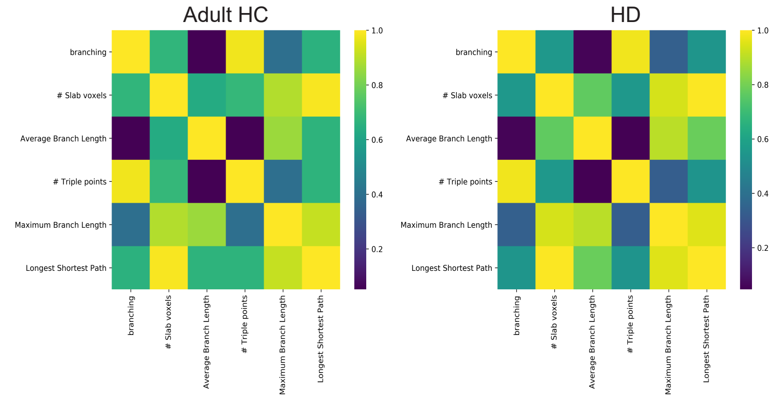

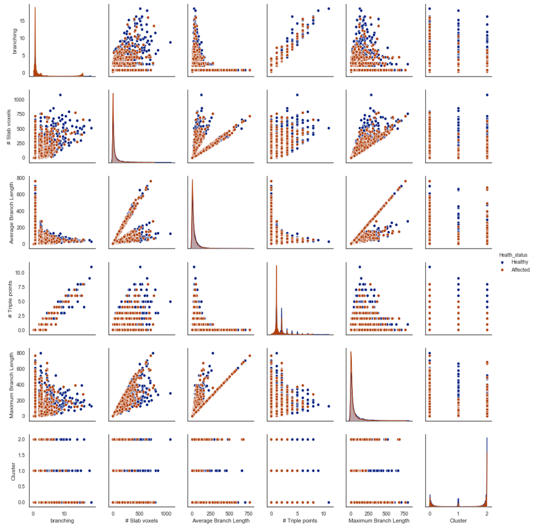

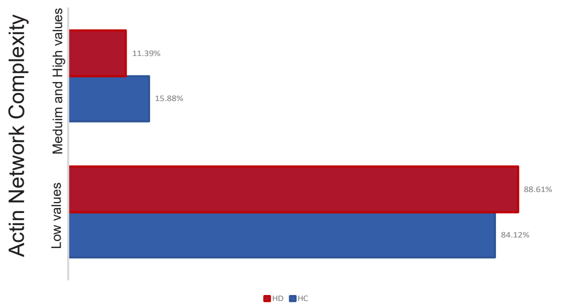

### Supplementary 5

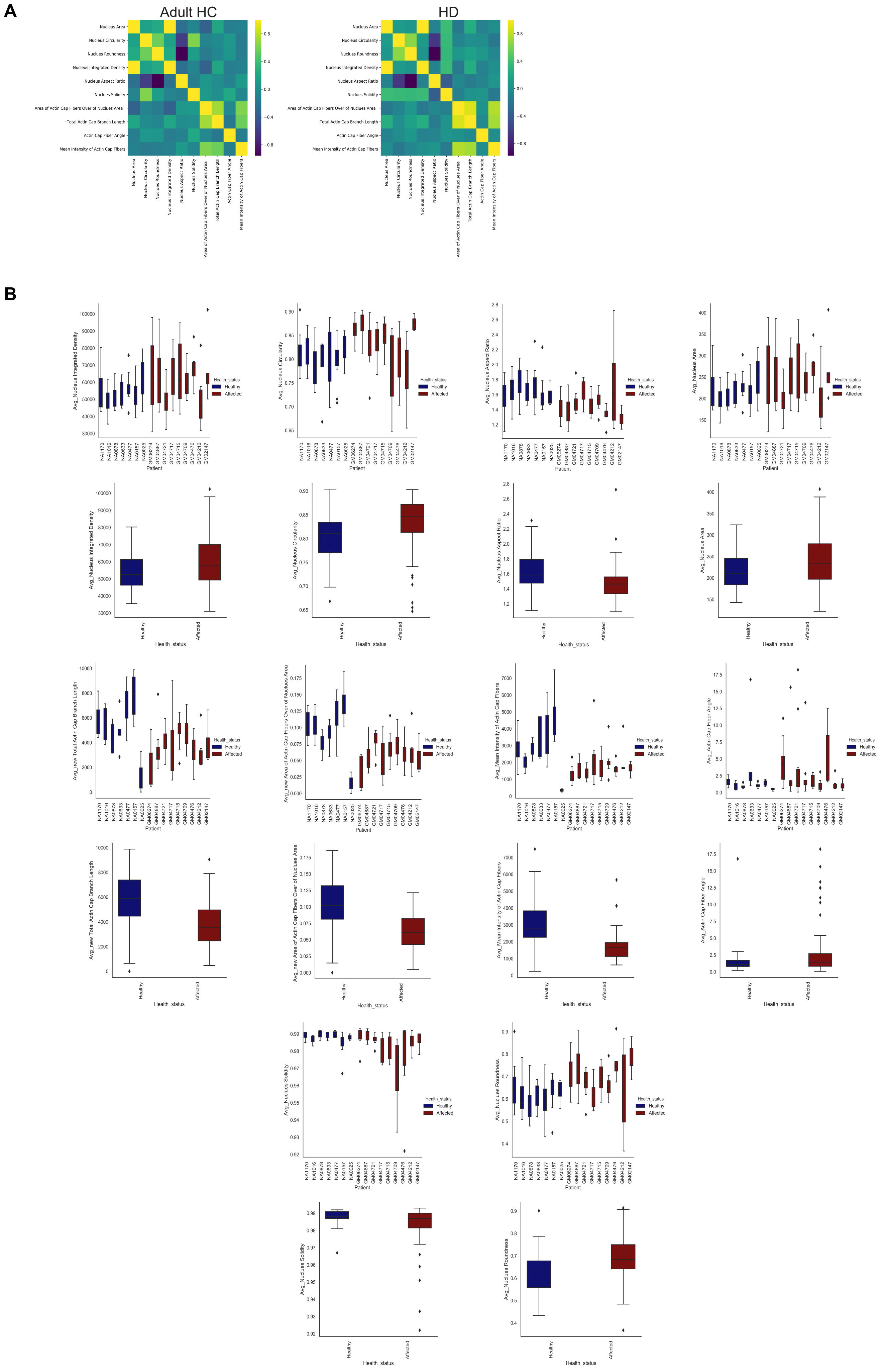

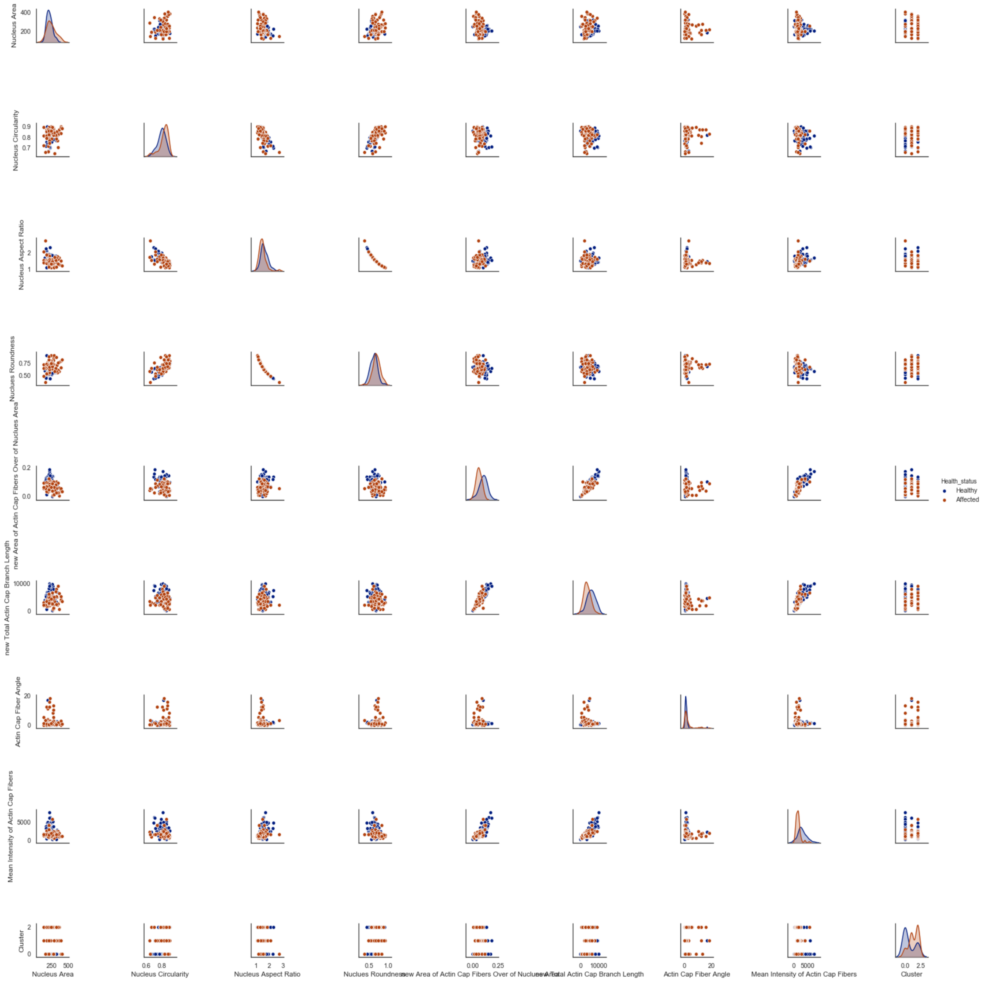

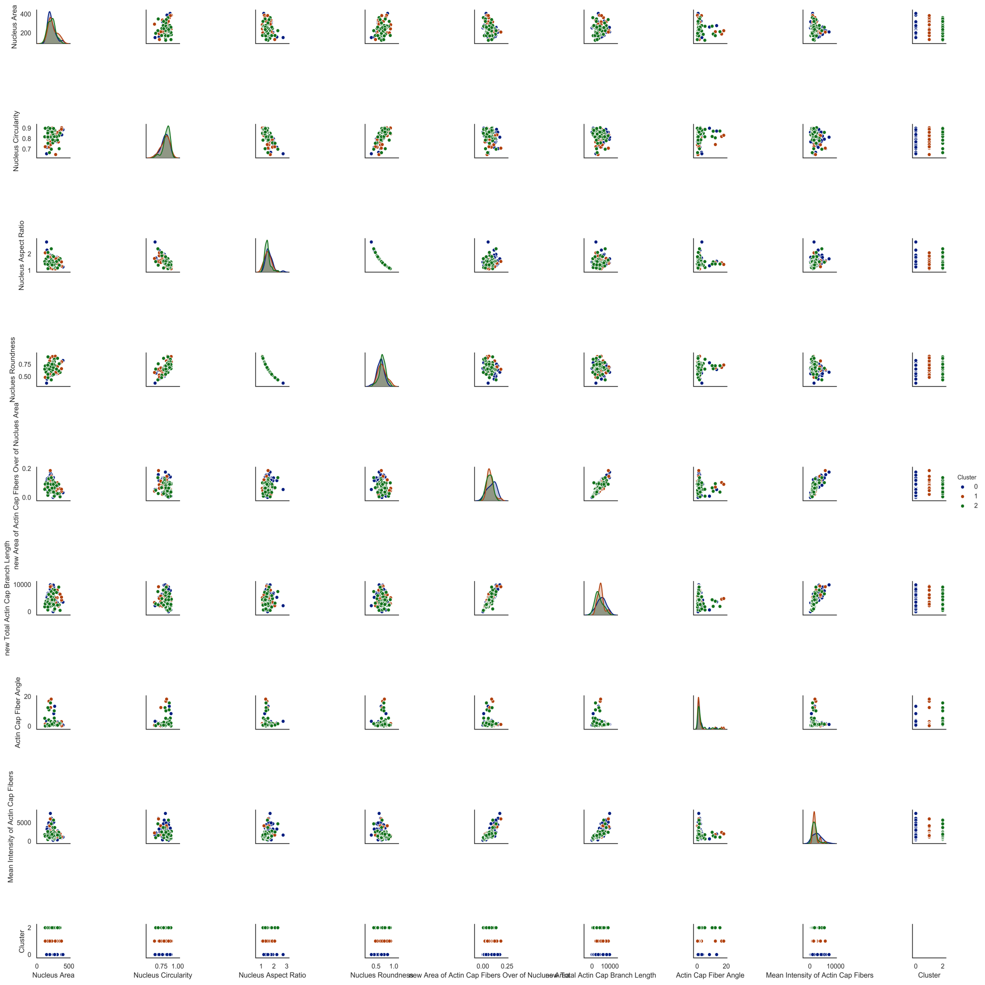
